# Substrate structure and computation guided engineering of a Lipase for Omega-3 fatty acid selectivity

**DOI:** 10.1101/592063

**Authors:** Tushar Ranjan Moharana, Nalam Madhusudhana Rao

## Abstract

Optimum health benefits of omega-3 fatty acids (ω-3 FAs) require it to be concentrated in its natural sources. Fatty acid selectivity of lipase governs the efficacy of the production of lipase-mediated ω-3 FAs concentrates. We attempted to improve the fatty acid selectivity of a lipase from thermophilic bacterium *Geobacillus thermoleovorans* (GTL) by two approaches. In a semi-rational approach, six amino acid positions of GTL interacting with the substrate, were identified by docking and were subjected to site-saturation mutagenesis. Three best substitutions were incorporated into GTL(CM-GTL). Hydrolysis of oil by lipase was monitored in a pH-Stat and the fatty acids released at various time points were analyzed by GC-MS.CM-GTL showed a significant improvement in discrimination against DHA during hydrolysis. In the second approach based on rational design, the active site was narrowed by incorporating heavier amino acids in the lining of acyl-binding pocket to hinder access to bulky ω-3 FAs. For this purpose, two amino acids surrounding the opening of the acyl pocket were replaced with the next heavier amino acids and the affinities were evaluated *in silico.* The double mutant, thus deigned, was found to be excellent in discriminating the ω-3 FAs during hydrolysis of triglycerides. Engineering the binding pocket of a complex substrate, such as a triglyceride, with the supportive information on substrate structure and its binding modes with the enzyme provided by computational methods, has resulted in designing two efficient lipase variants with improved substrate selectivity.

## Introduction

Health benefits of ω-3 fatty acids (FAs) as food supplements have been documented extensively [1-3]. ω-3 FAs helps in managing cardiovascular disease, stroke, obesity, arteriosclerosis, hypertriglyceridemia, inflammation and autoimmune diseases[4-10]. Their role in neuronal development is also well established[11]. American Heart Association recommends that patients with coronary heart diseases should consume one gram of ω-3 FAs daily. FDA has approved the use of ω-3 FAs concentrate as a drug against cardiovascular disease and hypertriglyceridemia. Many plant and fish sources, including Flax, Tuna and Anchovy contain substantial amounts of ω-3 FAs (∼30 mole %). cis-5,8,11,15,17-eicosapentaenoic acid (EPA) and cis-4,7,10,13,16,19-docosahexaenoic acid (DHA) are the major bioactive ω-3 FAs. To be used in pharmaceutical applications, the content of ω-3 FAs in oils should be more than 60 mole %. Studies revealed that health benefits depend on the omega-3/omega-6 ratio in oils and not merely on the omega-3 content[12,13].

One solution to the above problem is in concentrating ω-3 FAs in their natural sources. Various chemical methods such as urea complexation, low temperature crystallization, distillation, supercritical fluid extraction and chromatographic separation have been developed in order to concentrate ω-3 FAs [14-18]. However, each of these processes have issues on either economy of the process or on the assurance of product quality [19]. Lipase-mediated selective hydrolysis can provide a safer and cheaper alternative for concentrating ω-3 FAs. For the process to become effective, it requires a lipase with excellent fatty acid selectivity. Efforts to isolate a fatty acid-selective lipase or to improve the natural lipases for fatty acid selectivity have failed to provide fruitful results [20,21]. In this work, we took a lipase from *Geobacillus thermoleaverans* (GTL) as a model protein and engineered it to enhance its fatty acid selectivity using semi-rational and rational approaches.

*Geobacillus thermoleaverans* is a thermoalkalophilic bacterium. The lipase (GTL) is a 43kDa lipase with a lid and is active on oil/water interfaces. Lipases homologous to GTL show maximum activity between pH 9 and 10 and in the temperature range of 55 to 60°C[22]. All of them bind to one calcium and one zinc atom which enhance their stability[23]. Their T_m_ is close to 75°C[24]. Excellent activity and stability makes them apt for industrial applications. Large area of contact with the substrate makes them an ideal choice for fatty acid selectivity and provides options to alter their substrate preferences. However, their fatty acid specificity during hydrolysis has not been evaluated. We have evaluated the fatty acid selectivity of GTL and attempted to improve its fatty acid selectivity by protein engineering.

Both rational and semi-rational approaches have been very useful in altering substrate selectivity of enzymes[25-27]. Amino acids near the active site, which interact with the substrate during catalysis, play a major role in determining activity and fatty acid selectivity[28]. In the semi-rational approach, we have identified the amino acids that interact with the substrates during hydrolysis by covalent docking of substrate with the lipase using AutoDockVina[29]. We excluded the functionally important amino acids, and subjected amino acids which frequently interact with the substrate to site saturation mutagenesis (SSM). The best mutations were combined to test their substrate selectivity. In the rational approach, we exploited the structural difference between ω-3 FAs and non-ω-3 FAs. We hypothesized that by narrowing the fatty acid-binding channel in the active site of GTL, the binding of bulkier ω-3 FAs would be restricted and discriminated during hydrolysis. The binding energy differences were estimated by AutoDockVina and found to be favorable. The lipase variants thus generated with two amino acid substitutions were tested for fatty acid selectivity in anchovy oil. They showed excellent fatty acid selectivity and were able to concentrate ω-3 FAs in glyceride portion better than the GTL. Stability and activity of both the mutants were evaluated.

## Materials

Bleached anchovy oil was obtained from Ocean Nutrition Canada, Canada. Gum arabic, glycerol tributyrate, para-nitrophenyl butyrate (pNPB), hydrogen chloride-methanol solution, butylated hydroxytoluene (BHT) and methyl nonadecanoate were purchased from Sigma Aldrich, India. Silica gel TLC plates were purchased from MERCK, India. All the solvent and other chemicals used were of analytical grade or better. *Geobacillus thermoleovorans* strain (Acc. No. 4219) was purchased from MTCC, Chandigarh, India.

## Methods

### Cloning and expression of GTL

GTL gene was amplified from genomic DNA of *Geobacillus thermoleovorans,* signal peptide was removed and the resulting gene was cloned in to pET21d between Nco I and Hind III[30]. Mutations, identified by methods described below, were incorporated into GTL gene by overlap extension PCR method[31]. PCR mix was prepared by adding 20 ng pET21d plasmid containing GTL gene, 5 μl of 5 pm primer each to 25 μl Phusion master mix (2X) and diluting to 50 μl with MilliQ water. PCR was carried out for 30 cycles. Each PCR cycle consists of three steps: 1) denaturation at 98°C for 10 seconds 2) annealing at 55°C for 30 seconds 3) extension at 72°Cfor 30seconds/kb. In addition, 30 seconds of denaturation at 98°C at the beginning and 10 minutes of extension at 72°C at the end of PCR was carried out. First, fragments were generated by amplifying the first half of the gene with T_7_ forward primer and reverse sequence of appropriately designed primers. Similarly, the second half of the sequence was generated by designed primer and T_7_ reverse primer by PCR. The complete mutated gene was obtained by overlap extension PCR using the above fragments [31]. For Site Saturation Mutagenesis (SSM), MNN/NNK degenerate codon sets were used[32]. SSM was performed for each of the positions identified by docking. Sequences of the primers used in this study are provided in supplementary material (Supplementary Table 1).

### Modelling protein structures

All the protein structures were modelled by homology modelling using Modeller 9.14.[33]. PyMod 1.0 and PyMod 2.0, Pymol plug-ins were helpful in running modeller as well as Ramachandran plot analysis[34,35]. Open lid conformation of lipase L1 from *Geobacillus thermocatenulatus* (PDB ID 2w22, 95% sequence identity) was used as template[36].The sequence of the protein to be modelled was aligned with that of the template with the help of SALIGN module provided with MODELLER package. For modelling each protein structure, 10 structures were generated and the structure with the best DOPE score was selected[37]. Heteroatoms and water molecules were removed and additional energy minimization was avoided as these lead to poor models (data not shown).The generated model was validated by Ramachandran plot analysis using PyMod as well as analysis of other geometry parameters (bonds, angles, rotamers etc.; data not shown) using MolProbity[38].

### Semi-rational approach to design GTL for fatty acid selectivity

Amino acids which interact with the substrate play a critical role in its selectivity. In the semi-rational approach, we have identified the amino acids of GTL that interact with the substrate, i.e., triglyceride, by covalent docking of two representative substrates with GTL. Representative substrates were chosen based on the fatty acid composition and distribution among the three positions on glycerol backbone. Substrate 1 has EPA at sn-1, octadecanoic acid at sn-2 and 9-octadecenoic acid at sn-3 and substrate 2 has octadecanoic acid at sn-1, DHA at sn-2 and 9-octadecenoic acid at sn-3. AutoDockVina was used for docking calculation [29]. AutoDockTools and Pymol were used for the preparation of input files as well as data analysis [39,40]. Covalent docking was performed by flexible side chain method using a hydrogen atom as pseudo ligand[41]. Carbonyl carbon of the fatty acid undergoing hydrolysis was covalently attached to γ-oxygen of the active serine (S113) by Pymol. This structure was then processed by AutoDock Tool (MGL software package, version 1.5.6) where autodock atom type and charges were added. Enzyme without S113 side chain was saved as rigid receptor and S113 side chain covalently attached to triglyceride was saved as flexible residue. Kollman charges were added to the rigid part of the receptor, while Gasteiger charges were computed for the flexible side chain (triglyceride substrate covalently linked to S113)[42,43]. All the bonds, except double bonds and terminal bonds, were kept rotatable. Position and dimensions of grid box were such that it covers the entire active site and allow complete stretching of the flexible side chains. Grid spacing was kept at 0.375 Å. The 10 best binding modes for each substrate were analyzed to find the amino acids which interact with the substrate during hydrolysis. Amino acids that were frequently observed (at least 15 out of 20 modes) to be interacting with the substrates were considered to be important for substrate selectivity (Figure 1 and Table 1). However, many of the substrate-interacting amino acids play an important role in the activity of the enzyme and hence these positions were excluded and the remaining positions are considered for site saturation mutagenesis.

**Figure-1:**
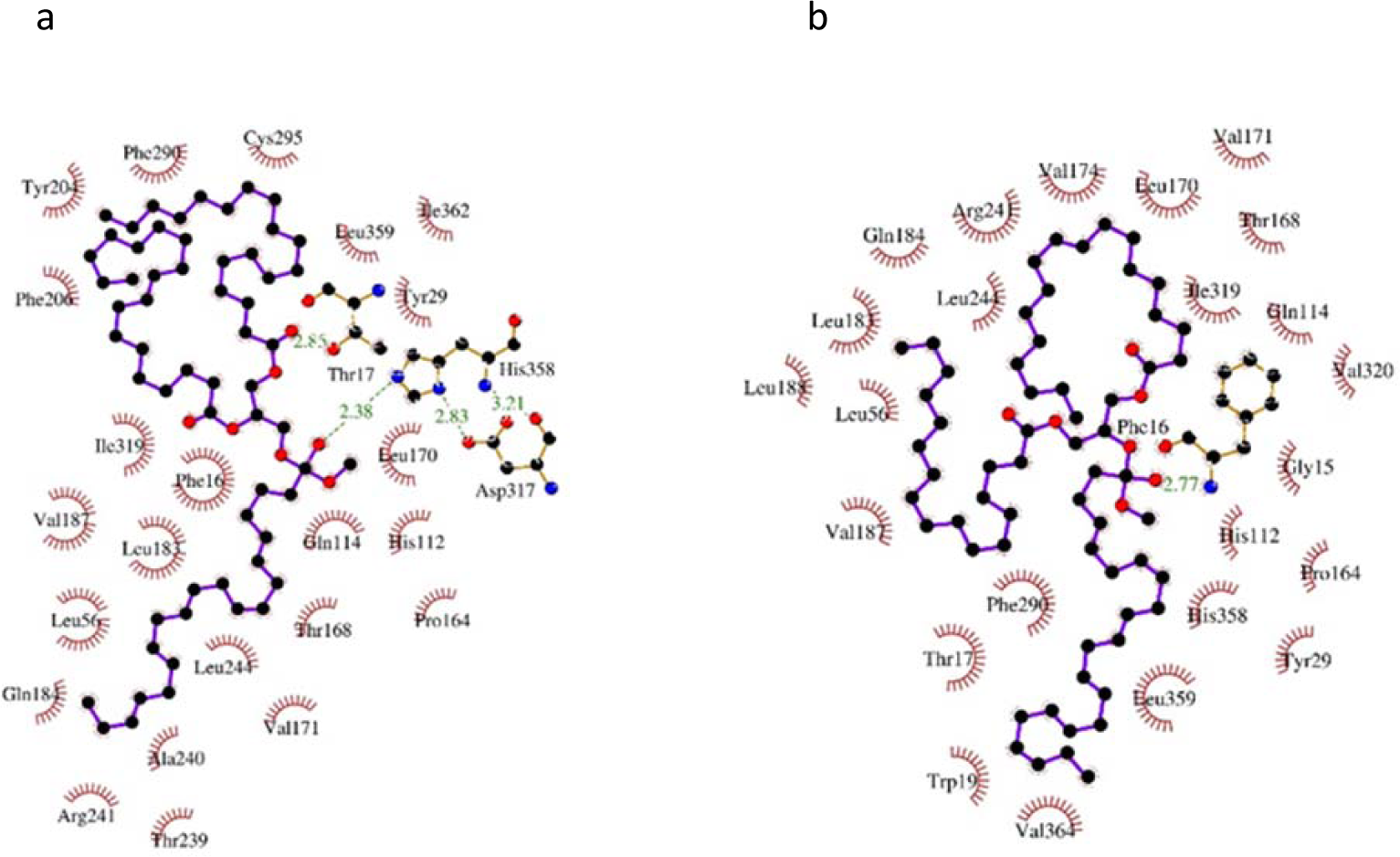
Amino acids interacting with triglyceride during hydrolysis as observed by covalent docking of representative substrates with GTL. Representative Substrate 1 has EPA at sn-1 position (a) and Representative Substrate 2 has DHA at sn-2 position (b) Other positions are occupied by octadecanoic acid and 9-octadecenoic acid. Docking was performed using AutoDockVina and the figure was generated using Ligplot[51].

**Table 1:** Interaction of substrate with GTL Amino acid positions which interact with the substrate during hydrolysis of triglyceride as predicted by covalent docking using AutoDock Vina.

| Frequency of interactions | Amino acid positions |
| --- | --- |
| 20 | 16, 17, 29, 112, 113, 114, 168, 241, 244, 319, 358 |
| 19 | 56, 183 |
| 18 | 171 |
| 17 | 170, 359 |

### Mutant library preparation

Amplicons of SSM, generated by the method described above, were cloned into pET 22b vector for periplasmic expression, between Nco I and Hind III restriction sites and transformed into *E. coli* BL21 DE3 expression system. Individual colonies were inoculated into a well of 96-well plate containing 200 μl of LBA (LB broth containing1mM ampicillin) and grown overnight at 37°C and at 180 RPM. Supernatants were removed after cold centrifugation at 1700 RCF for 5 min. *E. coli* cells in each well were resuspended in 100 μl of LBA. 100 μl of 60% sterile glycerol was added to each well and the plate was stored at −20°C.

### Screening the mutant library

To identify positive mutants, a two-stage medium throughput screening method was followed. The first stage is a qualitative assay which involves placing the culture supernatants containing mutant lipase in wells made in a tributyrin-agar plate. Lipase in the supernatants hydrolyzes tributyrin breaking the gum arabic-tributyrin emulsion, which leads to formation of transparent halo in a milky white tributyrin-agar plate[44]. This step was intended to screen out the inactive mutants. In the second stage, the active mutants identified in the first stage, were tested for their substrate selectivity by comparing their activity against two oils. Anchovy oil (rich in ω-3 FAs) and coconut oil (lacks ω-3 FAs) were chosen for this purpose. Using a pH indicator, phenol red, the activity of mutants was measured on a 96 well plate.

### Stage 1: Halo formation assay

20 μl inoculum from each well of the library was re-cultured in a corresponding well of a 96-well plate containing 200 μl LBA in each well overnight as described above. Cells were pelleted by cold centrifugation at 1700 RCF. 5 μl of the supernatant was placed in each well of tributyrin-agar plate and incubated at 37°C for overnight for halo formation.

Tributyrin-agar plates were prepared by mixing 7.2 ml of 0.5 M glycine (pH 9.5), 1.44 ml of 2.5 M NaCl, 0.72 ml of 0.5 M CaCl_2_ and 1.5 g gum arabic till gum arabic dissolved. To this mixture, glycerol-tributyrate (1.5 ml) was added and the solution was sonicated on ice to form a milky white emulsion. 0.4 g of agarose was dissolved in 40 ml of water by boiling and cooled to 40-50°C. The above emulsion was mixed with agarose and poured into a 150 mm diameter Petri plate. After the agar solidified, holes were made in 96 well plate format using gel punching apparatus.

### Stage 2: Differential activity assay

Active mutants from stage 1 were tested for their activity on anchovy and coconut oils. Differential activity assay was performed by measuring the activity of each of the mutant lipases on coconut oil (lacks ω-3 FAs) and anchovy oil (rich in ω-3 FAs) in a medium throughput manner. Buffer containing 25 mM glycine, 25 mM CaCl_2_, 1% gum arabic and 1% oil at pH 9.5 was emulsified by sonication. Phenol red (pH indicator) was added to a final concentration of 0.04 mg/ml. 200 μl of the above mixture was dispensed into each well of the 96-well plate and 20 μl of culture supernatant (obtained in the same manner as in Stage 1) was added and incubated in an incubator shaker at 37°C and 180 RPM. Pictures of the plates were taken at various time points to record the colour change. Based on the extent of colour change brought about by the mutants in comparison with GTL, the mutants were scored on a scale of 1 to 5, where a score of 1 indicates the least active mutant and 5, the most active mutant. The difference in scores obtained by a given mutant, with coconut oil and anchovy oil as substrates, was considered to evaluate the fatty acid selectivity.

### Rational approach to design GTL for fatty acid selectivity

ω-3 FAs are structurally very similar to saturated and monounsaturated fatty acids. Lack of polar atoms and highly flexible nature of the acyl chains reduces the structural diversity that is necessary for lipases to discriminate between the different types of fatty acids. Therefore, most of the natural lipases possess little fatty acid selectivity. The only structural difference between ω-3 FAs and non-ω-3 FAs is their bulkiness. ω-3 FAs, being *cis*-polyunsaturated, are bulkier compared to their saturated and monounsaturated counterparts. Few lipases discriminate between ω-3 FAs and non-ω-3 FAs to some extent based on their bulkiness [45]. Bulkier ω-3 FAs are poorly accommodated in the active site due to steric hindrance and consequently poorly hydrolyzed [46,47]. It is known that, out of the 4 pockets (namely HA, HB, HH and oxyanion hole) in the active site of *Geobacillus thermocatenulatus,*a close homolog of GTL, the HA pocket is ideally oriented with respect to catalytic Ser113 and it is lined by F16,S57, V171, V174,L183, V187,L188, L208, W211, L213,L244. It was observed that the fatty acid bound in the HA is subjected to hydrolysis and, henceforth, is referred to as acyl pocket (also referred to acyl binding site) [36]. In other lipases, it is known that selectivity of the lipase towards different substrates depends on the interaction of substrates with the acyl pockets [48,49]. A bent in the middle provides the characteristic “L” shape to the HA. This bent, which is also the narrowest part, was surrounded by amino acid F16, V171 and L183 (Figure 2). Excluding F16 which has a critical role in lid opening, V171 and L183 were replaced by the next bulkier amino acids i.e. leucine and phenylalanine, respectively. The GTL variant thus produced is referred to as DM-GTL.

**Figure-2:**
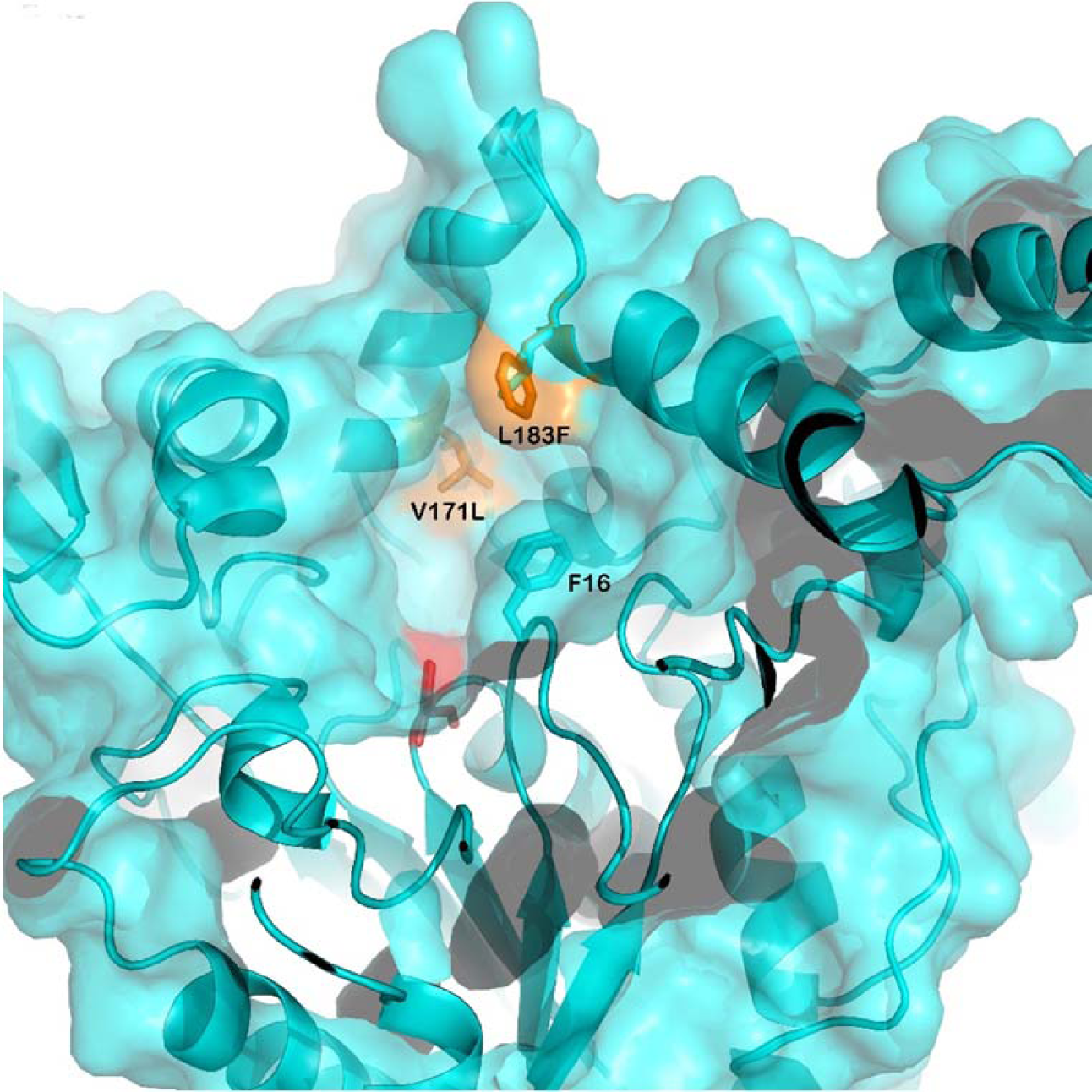
Mutation designed to narrow down the acyl pocket opening. Modelled structures of GTL and DM-GTL are superimposed and shown in cyan. Mutations are shown in yellow and active site serine is shown in red.

### *In silico* fatty acid selectivity of GTL and DM-GTL

*In silico* fatty acid selectivities of GTL and DM-GTL were estimated by calculating their affinities towards the representative substrates 1 and 2, while carboxyl carbon of different fatty acids was bound to γ-oxygen of S113, as employed in semi-rational approach. However, covalent docking using flexible side chain method does not provide interaction energy between receptor and covalently attached substrate. To obtain the interaction energies, a two-step method was used. In the first step, we calculated the best substrate binding mode. Only conformations, where the fatty acid which is undergoing hydrolysis (covalently attached to S113), occupies the acyl pocket were considered relevant. In the next step, we saved the coordinates (except Cα of 113SER) of the best binding modes of the flexible side chain and treated it as a ligand to calculate binding energy, without performing any further search (by activating option – score only).

### Overexpression and purification of lipases

Lipase genes, cloned into pET21D vector for cytoplasmic expression were transformed in to *E. coli* BL21 DE3 expression system. Cells were cultured in LBA till OD_600_ reached 0.6. Protein overexpression was induced by adding 1mM IPTG. After 3 hours of expression, cells were harvested by centrifuging at 4000 RCF. The cells were lysed by lysozyme treatment followed by sonication. Cell debris was removed by cold centrifugation at 10000 RCF. Lipases were purified by ethylene glycol gradient (0-80%) on an HIC (Hydrophobic Interaction Chromatography) column with phenyl Sepharose as stationary phase. Purity of GTL was confirmed by SDS-PAGE electrophoresis and the protein content was estimated by Lowry’ method using albumin standards.

### pNPB hydrolysis

4.2 mg of pNPB was dissolved in 1 ml of acetonitrile (20 mM). 20 μl of the above solution was diluted with 50 mM sodium phosphate at pH 7.5 to 1 ml. 0.1 μg of enzyme was added and the absorbance change was measured at 405 nm for two minutes to calculate the rate of hydrolysis.

### Hydrolysis of anchovy oil

Substrate for lipase was prepared by mixing 10 mM glycine, 10 mM CaCl_2,_ 5% gum arabic and 5% anchovy oil at pH 9.5, and the mixture was emulsified by sonication. Hydrolysis was initiated by adding 100 µg purified lipase at 37°C. Maintaining pH as well as monitoring reaction rate were carried out by titrating the released fatty acids against 1 M NaOH by a pH stat (Ω Metrohm 718 STAT Tritino). Samples were drawn at regular intervals. Free fatty acids were separated from glycerides by solvent extraction procedure. In this procedure, the mixture of free fatty acid and glyceride was mixed with 30% ethanolic KOH (0.5M) where free fatty acids get solubilised as potassium salt and insoluble glycerides were extracted twice with hexane. Free fatty acids were regenerated by lowering the pH below 2 with HCl [50]. The purity of separated fatty acids, as well as glycerides, was tested on silica gel TLC using hexane: diethyl ether: acetic acid (60:40:1, v/v) as mobile phase.

### Methylation and fatty acid analysis by GC-MS

Methylation of both free fatty acids was carried out by acid catalysis using hydrogen chloride–methanol as methylating reagent as described earlier[50], with one modification i.e., a commercial mixture of hydrogen chloride–methanol was used instead of preparing it by adding acetyl chloride in methanol. 1 μl of hexane containing fatty acid methyl esters (FAMEs) was analyzed by GC-MS (Agilent 6890). HP-5MS (30 mt X 250 μm X 0.25 μm) column was used for separation of FAMEs. Helium gas was used as carrier gas at a speed of 40 cm/sec. Injector temperature was kept at 250°C. Oven temperature was kept at 140°C for 2 min, increased to 290°C at a rate of 6°C/min and kept at 290°C for 3 min. The fatty acids were quantitated by using known quantities of an external fatty acid standard. Methyl nonadecanoate was used as an internal standard as described earlier [50].Selectivity for a given fatty acid (Fax) was calculated by using the following equation:

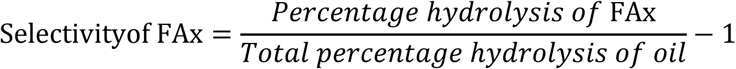

### Thermal unfolding of GTL mutants

Thermal melting of GTL, CM-GTL and DM-GTL was measured by recording the molar ellipticity at 222nm. Protein (0.1 mg/ml) in sodium phosphate buffer (10 mM, pH 7.5) was taken in a 1cm path length cuvette. Temperature was increased from 50°C to 90°C at a rate of 1°C/min using JASCO peltier type temperature controller and the ellipticity was measured using JASCO J-815 spectropolarimeter. Percentage unfolded fraction of protein was calculated using the following equation:

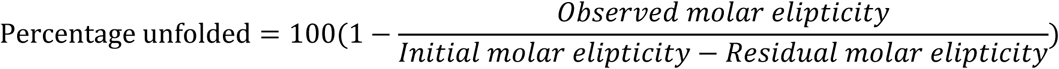

## Results

### Amino acid positions chosen for SSM

In order to know the amino acids that interact with substrate during hydrolysis, covalent docking of representative substrates with GTL was carried out. Figure 1 shows the amino acid interacting with representative substrates in the best mode of interaction. Best ten interaction modes for each substrate were considered. Amino acids which interact with substrate in at least 75% (15 out of total 20 modes) were considered as important for substrate selectivity (Table 1). However, many of the substrate interacting amino acids play an important role in the activity of the enzyme and hence these positions were excluded. Following amino acid positions were excluded from any substitutions for the stated reasons: F16 and T17 have critical role in structural rearrangement during lid opening; S113 is the active residue which carried out the hydrolysis; H112 stabilizes active serine (S113) in the closed conformation; Q114 and T168 form hydrogen bonds with substrate (data not shown); R241 forms salt bridge with D178 which stabilizes open lid conformation. After exclusion of these amino acid positions, amino acid positions 170, 171, 244, 319, 358 and 359 were chosen for SSM (Supplementary Figure 1).

### Screening active mutants

In the first stage of screening, we performed qualitative halo formation assay to identify active and inactive mutants using tributyrin as substrate. For each amino acid position at least 150 colonies were picked to ensure inclusion of all possible amino acid substitutions at each position [32]. Mutation at some positions had little impact on activity, while at other positions the substitutions resulted in mostly inactive enzyme (Figure 3). Out of the 960 individual mutants, generated by SSM at six positions, 210 colonies showed lipolytic activity and were subjected to a second screen

**Figure-3:**
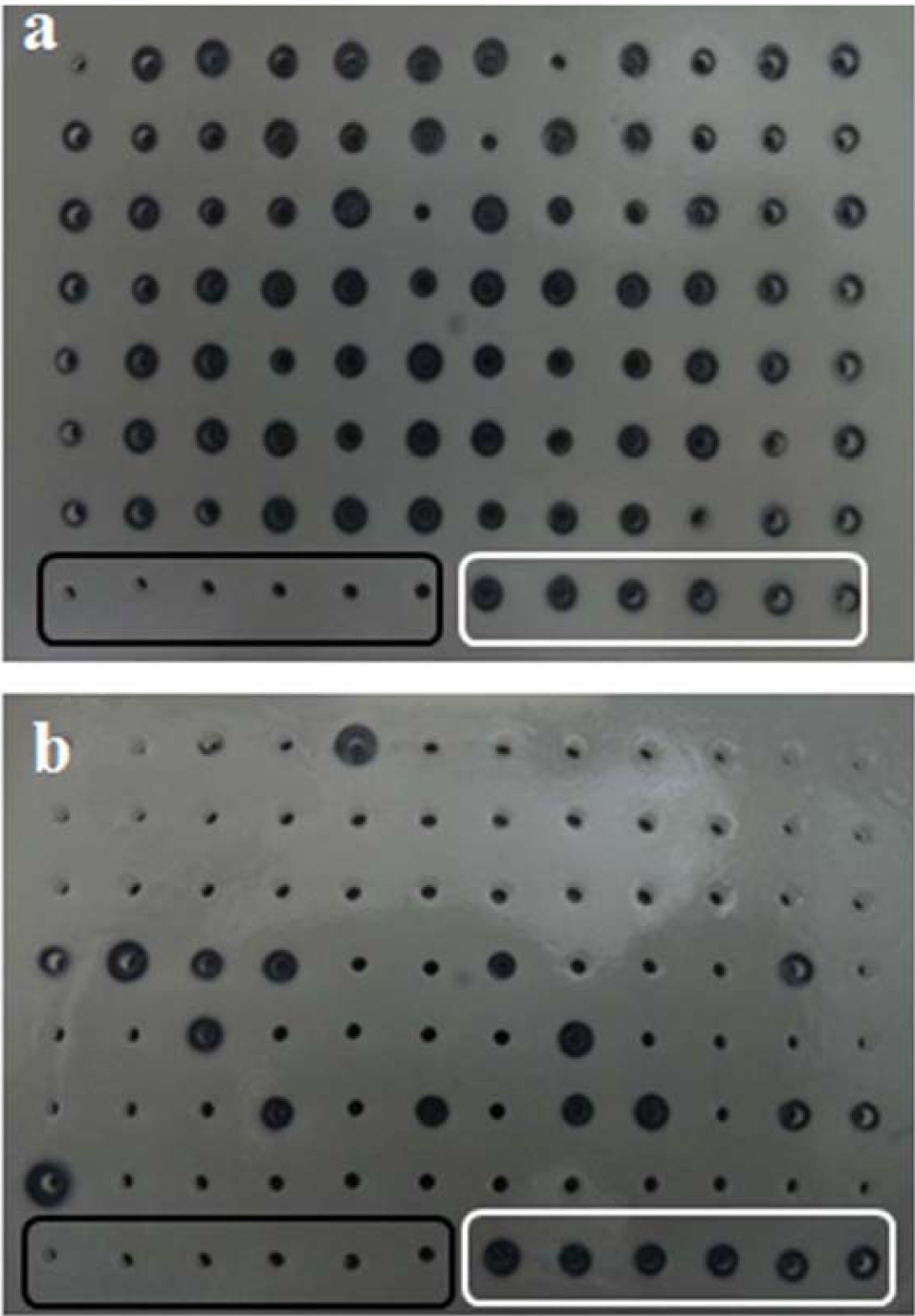
Halo formation by mutants of GTL on agar-tributyrin plate. Wells within black rectangle are negative controls (without any lipase) and wells within white rectangle are positive controls (contain GTL). Most of the mutants at position 244 are active (a), while mutants at position 359 are mostly inactive (b).

### Screening for differential activity

In the second stage of our screening, fatty acid selectivities of the 210 active mutants were evaluated by comparing their activities on two oil substrates with different fatty acid compositions. The oils used were coconut oil, which lacks ω-3 FAs and anchovy oil, which is rich in ω-3 FAs. Activity was measured in a medium throughput way, by recording the time required for decrease in pH due to release of fatty acids. Changes in pH were observed by the pH indicator dye, phenol red, which changes colour from red (above pH 7) to yellow (below pH 7). Figure 4 shows a representative plate that demonstrates the colour change due to hydrolysis of anchovy oil by GTL mutants.

**Figure-4:**
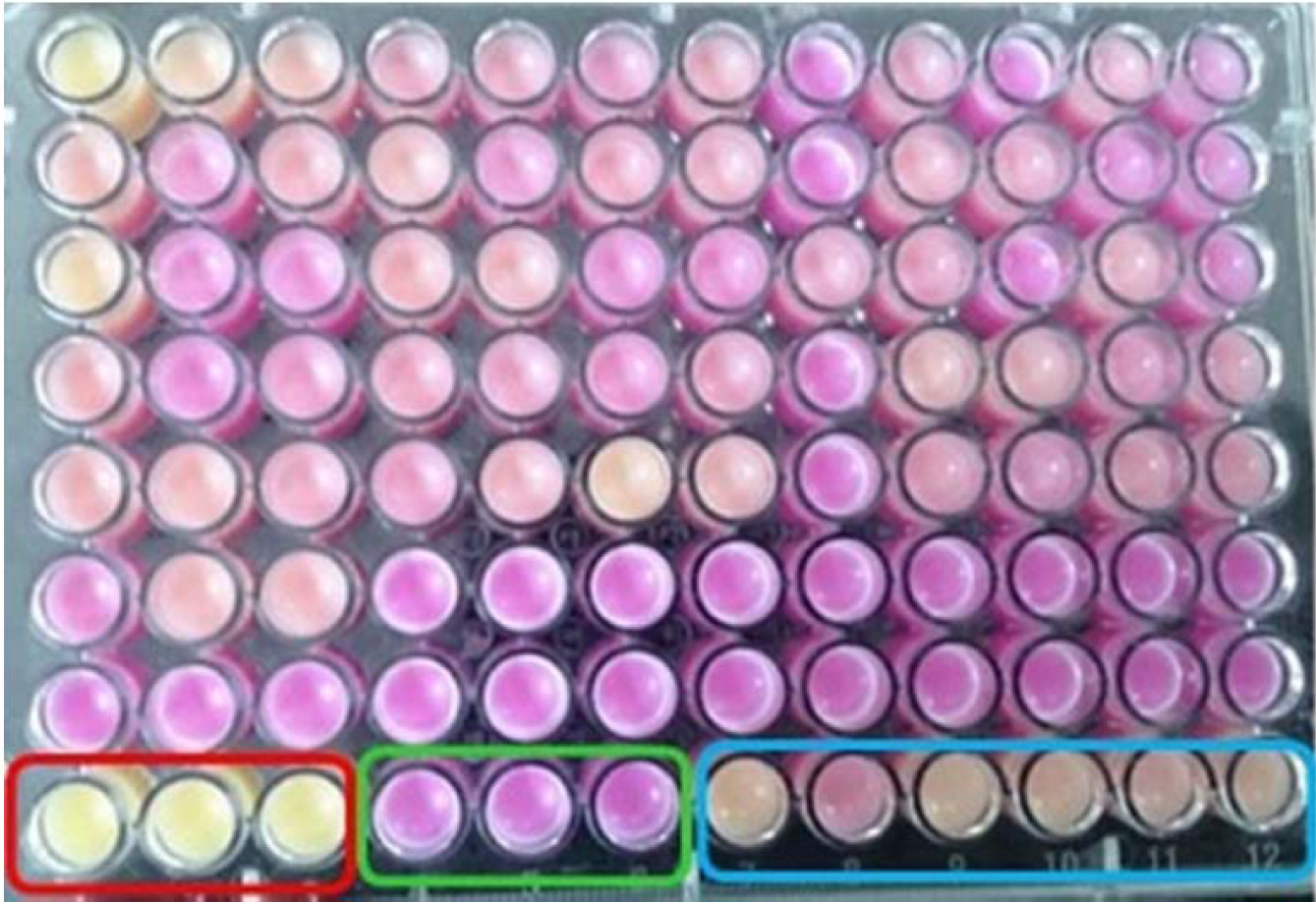
Change of pH due to hydrolysis of triglyceride by mutants of GTL indicated by phenol red. Wells in red rectangle are positive controls (pH <6, without lipase), in green rectangle are negative controls (pH 9.5, without any lipase) and in blue rectangle are with GTL. Acidic samples are yellow in colour (due to activity of lipase) and alkaline samples are purple in colour.

We hypothesized that the difference in activity in two oils by the mutants could be because of fatty acid selectivity. Since we already know that GTL preferentially hydrolyzes saturated fatty acids, mutants which showed comparatively more activity in coconut oil were considered positive and the corresponding lipase genes were sequenced. Out of the 210 mutant colonies screened, 28 colonies showed improved hydrolysis of coconut oil over anchovy oil compared to that of the GTL. Out of the 28 mutants sequenced, in 6 mutant sequences the base substitution did not result in amino acid substitution. The remaining 22 sequences were found to contain 13 different mutations. These were found at positions 170, 171 and 359 (Figure 5). At position 170, leucine was replaced by glycine in four lipase mutants and by tryptophan in two mutants. More diverse mutations were observed at position 171, where valine was replaced by arginines in three mutants, by proline in two, by tryptophan in two mutants and by alanine, phenylalanine, lysine and threonine in one mutant each. Mutations at position 359 include the replacement of leucine by cysteine in two mutants and by alanine, lysine and threonine in one mutant each. The most frequent mutations at each of these 3 positions are L170G, V171R and L359C. A lipase variant with these mutations was generated by SDM (named as CM-GTL). CM-GTL was cloned into expression vector, overexpressed and purified. The CM-GTL fatty acid selectivity was investigated along with the mutants generated by rational approaches as described below.

**Figure-5:**
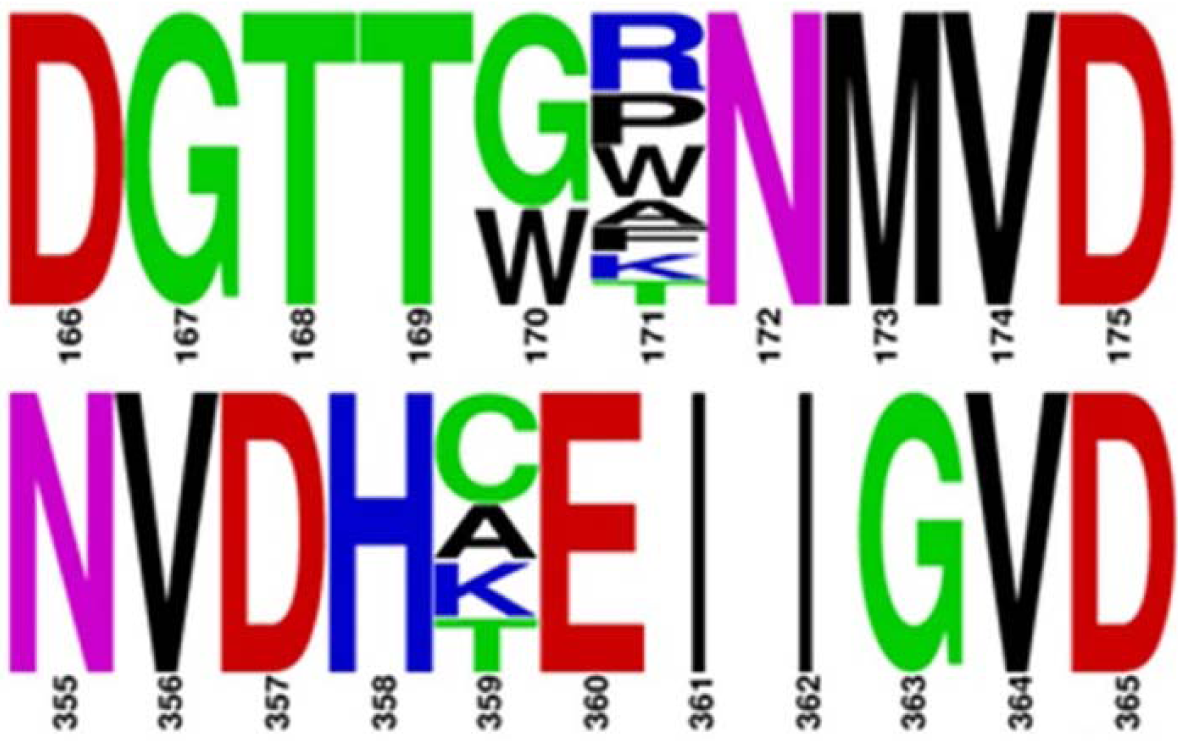
Mutations at different positions which showed improved differential activity during screening. Occurrences of GTL amino acids at the positions of mutation are ignored for better presentation. The figure and the colour code is generated by WebLogo[84].

### *In silico* fatty acid selectivity of GTL and DM-GTL

As described in the methods section on designing a lipase variant using rational approach, a double mutant, named as DM-GTL, was generated with following amino acid substitutions V171L and L183F. The positions were arrived after modelling the binding of substrate to various pockets in acyl chain binding in GTL. Table 2 summarizes the binding energies of both the triglycerides when each of the three fatty acids of the glycerides is in a configuration that is suitable for hydrolysis. It is clear that DM-GTL has less affinity towards EPA and DHA hydrolysis and more affinity for saturated fatty acids compared to that of GTL. DM-GTL showed a mixed effect on mono unsaturated fatty acids. The mutations were incorporated in GTL and the resulting mutant was tested for its fatty acid selectivity.

**Table 2:**
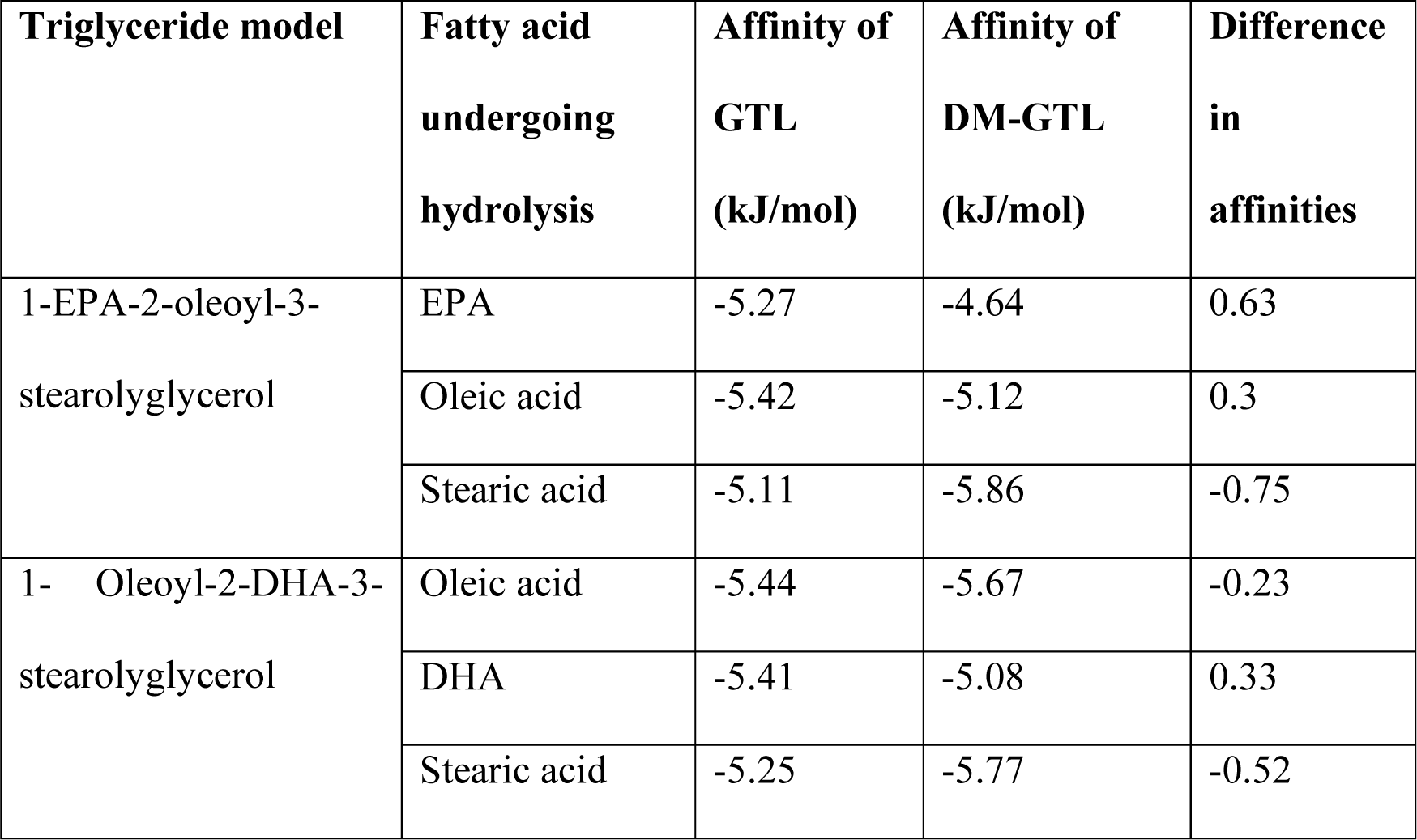
Affinities of two designed substrates with GTL and a mutant Affinities of GTL and DM-GTL towards hydrolyzing different fatty acids. Affinity of any fatty acid is calculated to be the binding energy between the representative substrate and enzyme when that fatty acid is undergoing hydrolysis.

### Fatty acid selectivity of GTL DM-GTL and CM-GTL

The rate of hydrolysis of anchovy oil by GTL, CM- and DM-GTL was monitored by pH-Stat. During hydrolysis, aliquots of the reaction were taken at various time points for processing to separate the fatty acids from the unhydrolyzed portions. Free fatty acids were methylated and the derived FAMEs were quantified using GC-MS. The data presented in Figure 6 shows the selectivity of the lipases at various extents of hydrolysis. Value of selectivity can vary between −1 to 1, where “-1” means complete discrimination (no hydrolysis) and 1 means complete preference (100% selective). All three lipases preferentially hydrolyzed saturated fatty acids at the beginning of the reaction and the selectivity parameter plateaued with increase in hydrolysis. Both CM-GTL and DM-GTL showed an improvement in selectivity compared to the GTL. Hydrolysis of individual fatty acids was estimated by GC-MS. The discrimination of EPA hydrolysis was unaltered between GTL and CM-GTL while the discrimination for DHA changed from −0.33 for GTL to −0.75 for CM-GTL (Figure 6b). This increase in discrimination in hydrolysis by CM-GTL was, although to a lesser extent, retained till the end of the reaction time. On the other hand, DM-GTL showed improvement in selectivity for both EPA (from −0.5 to −0.83) and DHA (−0.33 to −0.95). Also, DM-GTL retained this selectivity for a longer duration (Figure 6c). However, the ability to discriminate between different fatty acids during hydrolysis was compromised after the extent of hydrolysis crossed 30 percent. To obtain a quantitative comparison among the variants, discrimination (magnitude of selectivity) of GTL, CM- and DM-GTL towards EPA and DHA was compared after 20 percent hydrolysis (Supplementary Figure 2). The data shows excellent discrimination by DM-GTL for both EPA and DHA, while discrimination of CM-GTL increased only for DHA.

**Figure-6:**
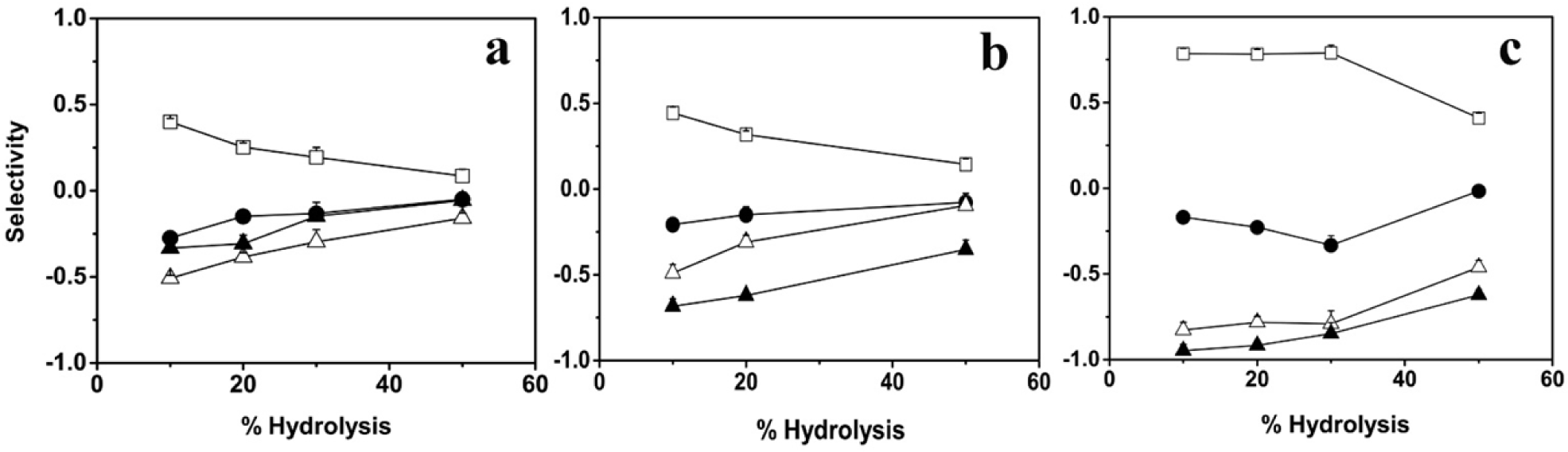
Activity of three lipases with anchovy oil as substrate. Activity monitored by a pH Stat method. GTL (dark circle), DM-GTL (dark square) and CM-GTL (open circle).

All the three lipases hydrolyzed saturated and mono unsaturated fatty acids more efficiently than ω-3 FAs which indicated that the ω-3 FAs are accumulated in the glyceride fraction. To evaluate their ability to concentrate ω-3 FAs, we compared EPA and DHA content of glyceride layer after 20 percent hydrolysis by GTL and DM-GTL, which shows that both the enzymes enriched ω-3 FAs (Supplementary Figure 3). While GTL showed enrichment of EPA only, DM-GTL showed significantly better enrichment in both EPA and DHA.

### Thermal stability of GTL mutants

Being an enzyme from a thermophile, GTL is highly thermostable. To assess the influence of the newly incorporated mutations on the thermal stability, GTL mutants were characterized. Thermal unfolding of GTL, CM-GTL and DM-GTL was observed by CD at 222 nm [52]. Figure 7 shows the unfolding of the three lipases upon heating. The mid-point of melting (T_m_) transition for GTL was estimated to be 76.5°C. The T_m_ of the mutants (CM-GTL and DM-GTL) was lower than that of GTL by 3°C, suggesting that the mutations have partially destabilised the native protein. The thermal transition profile of DM-GTL, like GTL, exhibits a sharp transition, indicating cooperativity, whereas the width of the thermal transition of CM-GTL was broader, indicating less cooperativity during unfolding of CM-GTL.

**Figure-7:**
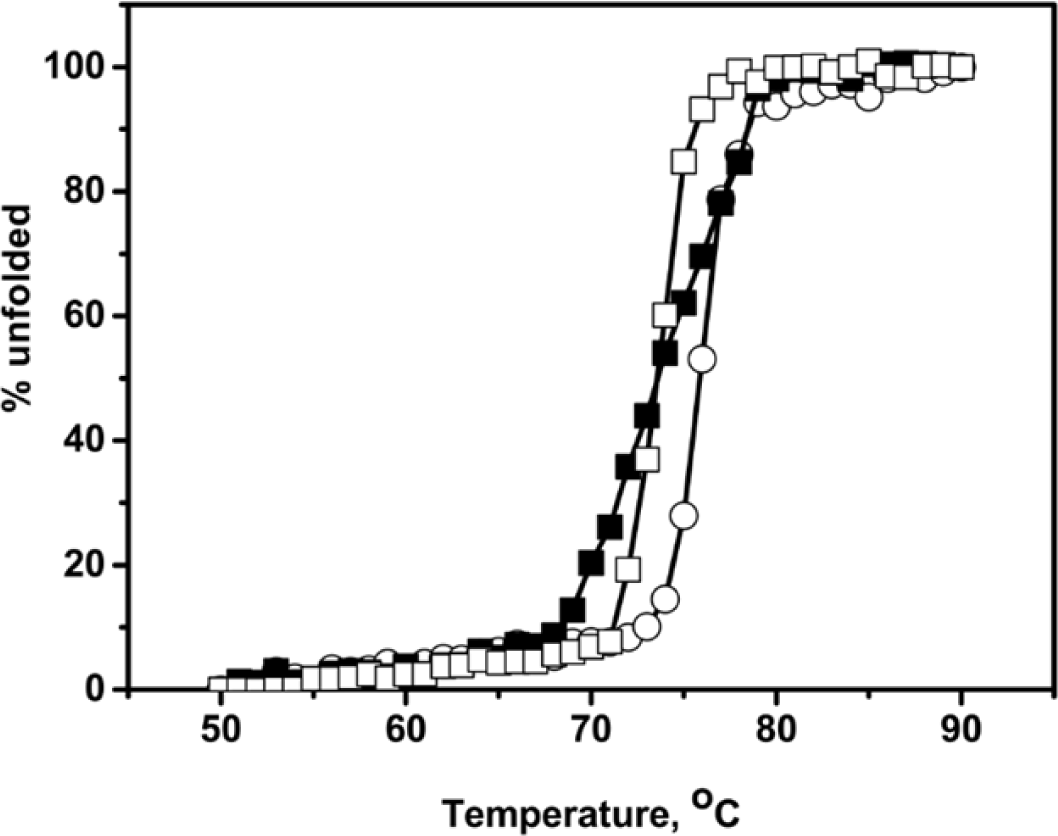
Thermal unfolding of GTL and its mutants. Fraction unfolded was calculated from the ellipticity at 222 nm. GTL is represented by circle, DM-GTL by open square and CM-GTL by dark square.

### Activity of GTL and its mutants

Mutation at the active site can affect the activity of enzyme. Since all the mutations incorporated are in the active site of the lipase, we investigated the activity of lipases on water soluble choromogenic substrate, pNPB, using spectrophotometer and on water insoluble anchovy oil by pH Stat method. Activity (unit = micromoles pNPB hydrolysed /min/mg of protein) of GTL, DM-GTL and CM-GTL was 161.5, 210.6 and 16.8 units respectively with pNPB as substrate. Compared to GTL, the activity of DM-GTL improved significantly whereas the activity of CM-GTL was severely compromised. With anchovy oil, the observed activities (unit = micromoles of OH added /min/mg of protein) estimated by pH Stat method, were 312.5, 187.5 and 37.5 units with GTL, DM-GTL and CM-GTL, respectively (Figure 8). The data suggests that the incorporated mutations reduced the activity of DM-GTL for anchovy oil but not for pNPB. CM-GTL activity was significantly reduced compared to GTL with both the substrates.

**Figure-8:**
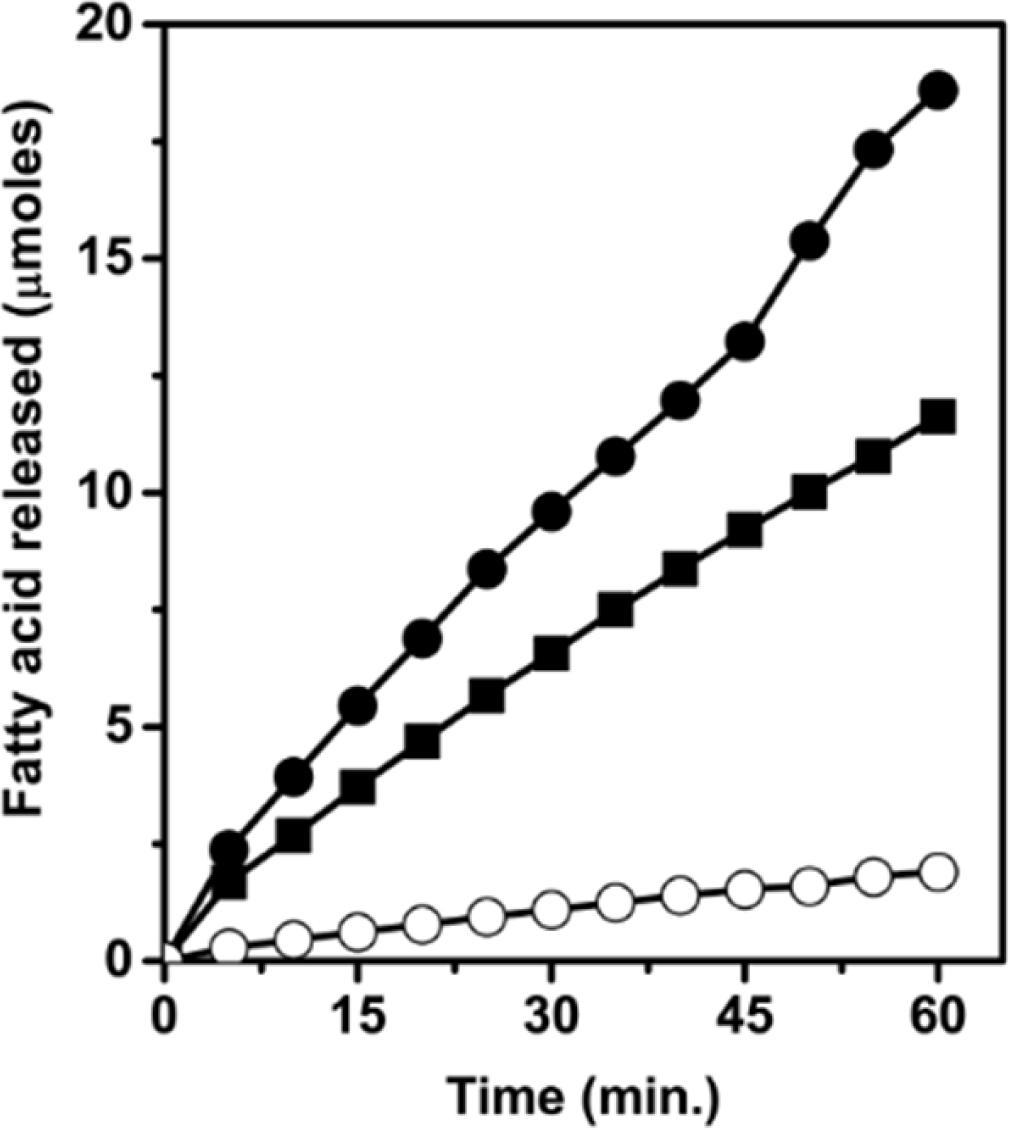
Fatty acid selectivity of GTL (a), CM-GTL (b) and DM-GTL (c) at various extents of hydrolysis of anchovy oil. Saturated fatty acids (Open square), mono-unsaturated (Dark circle), EPA (Open triangle) and DHA (Dark Triangle). Selectivity value 0 indicates neutral hydrolysis, 1 indicates total preference and −1 indicates total discrimination (no hydrolysis).

## Discussion

Among all the methods of concentrating ω-3 FAs in triglyceride form, selective hydrolysis of fatty acids by lipases is considered to produce a cleaner product and at less environmental load. Also, in this method mostly ω-3 FAs get concentrated in the form of triglycerides, which was shown to be biologically more valuable than free ω-3 FAs [50,53-55].Substrate selectivity of the lipase is the key factor for the concentration, which determines the yield and economics of the process. Since most of the known lipases have low fatty acid selectivity, it is essential to enhance their selectivity before they can be successfully used to concentrate ω-3 FAs. Several strategies, including substrate imprinting, enzyme immobilization, enzyme entrapment, reaction engineering, solvent engineering, protein engineering, have been used to enhance fatty acid selectivity of lipases with only a marginal success [56-61]. Though challenging, protein engineering is the most effective as well as most economical method to alter substrate preferences.

Protein engineering has been successfully used to alter substrate selectivity of many enzymes including several lipases [62-64]. *Candida rugosa* lipase has been engineered for selectivity of fatty acid chain length by blocking the active site tunnel[65]. *Candida antarctica* lipase B has been successfully engineered for enantio selectivity based on the enzyme-substrate interaction [64]. In this study, decreased volume of stereo-selective pocket correlates with increased stereo-selectivity. Enzyme selectivity also depends on the active site flexibility, wherein a less flexible site is more selective [66]. All these results are in line with our rational approach which results in DM-GTL with excellent fatty acid selectivity. The DM-GTL, constructed based on narrowing of the acyl binding pocket in the active site of GTL showed excellent fatty acid selectivity till 20 percent hydrolysis of anchovy oil and enriched ω-3 FAs in the glyceride fraction.

Selective hydrolysis of saturated fatty acids leads to increase in concentration of ω-3 FAs in the glyceride layer. At 20% hydrolysis concentration of ω-3 FAs increased from less than 30 mole% to over 40 mole%. Similar levels of enrichment, with best lipases have been observed after 50 to 80% hydrolysis [67]. In the studies where higher efficiencies in concentration of ω-3 FAs were observed, these efficiencies were shown to be achieved by repeated hydrolysis [68], trans-esterification with ω-3 FAs[69], or combining lipase catalysis with other methods [70,71]. Taiwo *et al.* have achieved up to 80% ω-3 FAs concentration in fish oil hydrolysate [72]. However, the reported concentration is the concentration of ω-3 FAs as mono-glyceride. The actual concentration in glyceride mixture will be many folds less as mono-glyceride constitutes a very small fraction of fish oil hydrolysate. Also, unlike other lipases which concentrate only one out of many ω-3 FAs, DM-GTL is effective in concentrating both EPA and DHA [71,73]. Selectivity of the lipase can be further improved by either narrowing the end of the acyl pocket, which might lead to improved distinction between ω-3 FAs and other unsaturated fatty acids or by narrowing the other acyl chain pockets, which may lead to distinction between ω-3 FAs and non-ω-3 FAs in alternate binding modes. Also ω-3 FAs are concentrated as DG and MG instead of TG. The obtained glyceride can be trans-esterified to obtained ω-3 FAs concentrate as TG [69].

The enhancement in discrimination against ω-3 FAs observed in this study by narrowing the binding channel requires structural proof to rule out other contributions of the mutations in altering the specificity. To confirm the effect of substitutions on the acyl binding pocket, we attempted a wide set of conditions to crystallize the DM-GTL. In some conditions, we obtained very small crystals that were not suitable for diffraction (data not shown). This study is in progress and we expect to get the crystal structure of the mutant.

Semi-rational approaches also delivered several success stories [74,75]. CM-GTL, constructed based on the SSM on the substrate contact sites of the active site amino acids in GTL, also showed improved selectivity, however, the improvement was not as significant as that of DM-GTL. At 20% hydrolysis we did not expect significant improvement in the enrichment of ω-3 FAs with CM-GTL compared to GTL, hence the product profile was not evaluated. All the mutations observed in the screening were concentrated into three positions, out of which two were in the acyl pocket. This observation is in line with the previous works where people have altered the substrate selectivity by incorporating mutations in the acyl pocket [76,77]. Mutations most frequently observed upon screening at each position were combined to enhance fatty acid selectivity. Examining all the combinations of mutations is not practical. Therefore, we limit our study to one mutant with most frequent substitutions, although the epistatic effect could be either synergistic or antagonistic.

We have also evaluated the effects of mutations on activity and stability of the lipase. Both CM- and DM-GTL showed poor activity on anchovy oil. The activity of DM-GTL on anchovy oil was 65% that of GTL and the activity of CM-GTL was 10% that of GTL activity. This result is as per our expectation as the mutations were incorporated to disfavour ω-3 fatty acids which constitute 30 percent of anchovy oil. DM-GTL hydrolyzed pNPB better than GTL. This could be because a greater fraction of DM-GTL was in the lid open conformation in the aqueous phase. Poor activity of CM-GTL on both the substrates could be because of the proximity of the L359C mutation to the active site than other mutations. The incorporation of mutations in the GTL have reduced the thermostability of the GTL to some extent; however, this decrease in stability is marginal and therefore does not hinder the usability of the lipase.

GTL as well as the CM- and DM-GTL mutants lose their selectivity with the progress of hydrolysis, which is probably due to buildup of the products in the form of mono-, diglycerides and free fatty acids. Similar phenomena have been observed for other lipases as well [78,79]. The continuously altering composition profile of the lipase “substrate” and consequent changes in the nature of the substrate emulsions lead to further loss of selectivity. The presence of alternative substrate binding mode in other lipases is also being speculated as a possible reason for the loss of selectivity [80]. If present, these can become a prominent mode of binding in later part of the hydrolysis and may compromise the selectivity. Altering fatty acid selectivity of lipases during hydrolysis of natural oil poses several formidable challenges. Highly flexible structure and lack of polar atoms in fatty acids makes it difficult for any enzyme to select one fatty acid over the other. Natural oil sources of ω-3 FAs, such as anchovy, Tuna etc. usually contains several species of triglycerides with varying fatty acid composition and also with variations in their positional distribution[81]. Hydrolysis of a triglyceride molecule may lead to six different possible products depending on which bond is cleaved. Some of the products (di and mono glyceride) also simultaneously can act as substrates confounding the kinetics. Also, fatty acid selectivities of many lipases depend on the overall substrate structures [82]. Compositional complexities of natural oils further increase the difficulty in arriving at a designing strategy. Lipases without lid, mostly do not show fatty acid selectivity. In these lipases, the area of contact between the active site and their substrates was not extensive enough to distinguish between different fatty acids. Lipases with lid, such as GTL, require interfacial activation. This requires emulsifying reagent and involves major structural rearrangement near the active site. Both the emulsifying reagent and the structural rearrangement affect substrate selectivity and are difficult to model[83].

## Conclusion

We have improved the fatty acid selectivity of GTL by both rational and semi-rational approaches. Lipase engineered by the rational approach (DM-GTL) showed almost complete discrimination against ω-3 FAs at the beginning of the hydrolysis. With the progress in hydrolysis, selectivity starts decreasing. Further improvement of selectivity on the designed mutants might open the door for economical enrichment of ω-3 FAs. While mutations lining acyl pocket are obvious to alter substrate selectivity, mutations at other positions can also contribute to it. The complexity of the substrate along with poor knowledge about alternate substrate binding modes are the challenges to overcome for rational designing of lipases.

## Acknowledgments

We acknowledge Indo-Australian Biotechnology Fund (GAP373) for the financial support and TRM acknowledges the research fellowship received from CSIR. The authors thank Drs. Colin Barrow and T Akanbi for gracious inputs during the initial stages of the work. The authors declare no conflicts of interest to disclose.

## Author contribution

TRM has performed the experiments and the simulations presented in the article. NMR has conceived the idea, organized and written the article.

