## Supplementary Material for "Substrate structure and computation guided engineering of a Lipase for Omega-3 fatty acid selectivity"

**CSIR-Centre for Cellular and Molecular Biology**
**Uppal Road**
**Hyderabad India 500007**

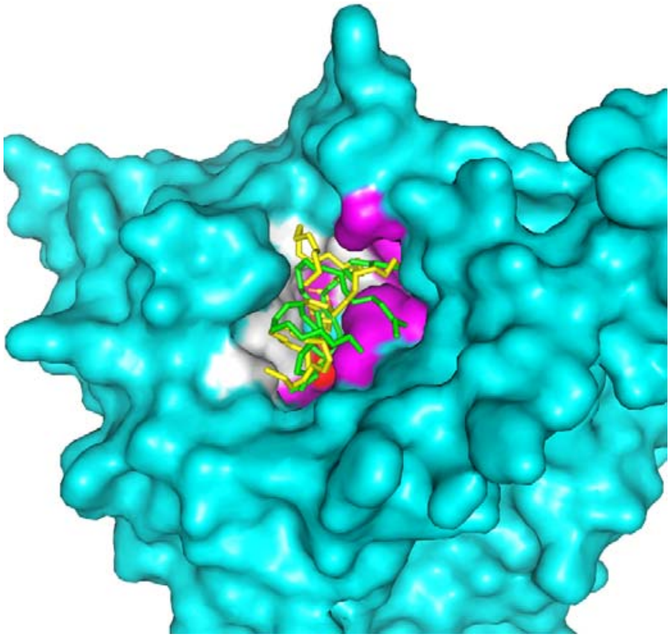

**Supplementary Figure 1:** Covalent docking of triglyceride with GTL. Amino acid positions chosen for SSM were shown in white, while amino acids interacting with substrate, but not included for SSM, are shown in magenta and active serine (S113) is shown in red. Triglyceride molecule shown in green represents substrate as shown in Fig.1a and the molecule shown in yellow represents substrate as shown in Fig.1b.

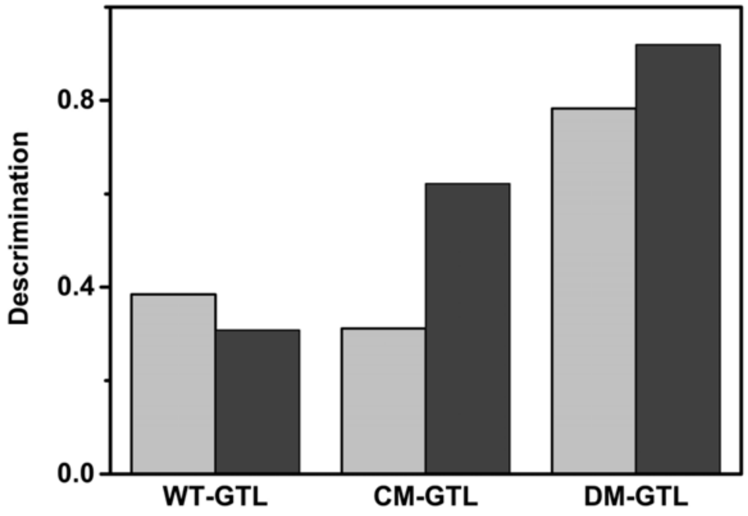


**Supplementary Figure-2:** Fatty acid discrimination of GTL, CM-GTL and DM-GTL at 20 percent hydrolysis. EPA (Light grey) and DHA (Dark).

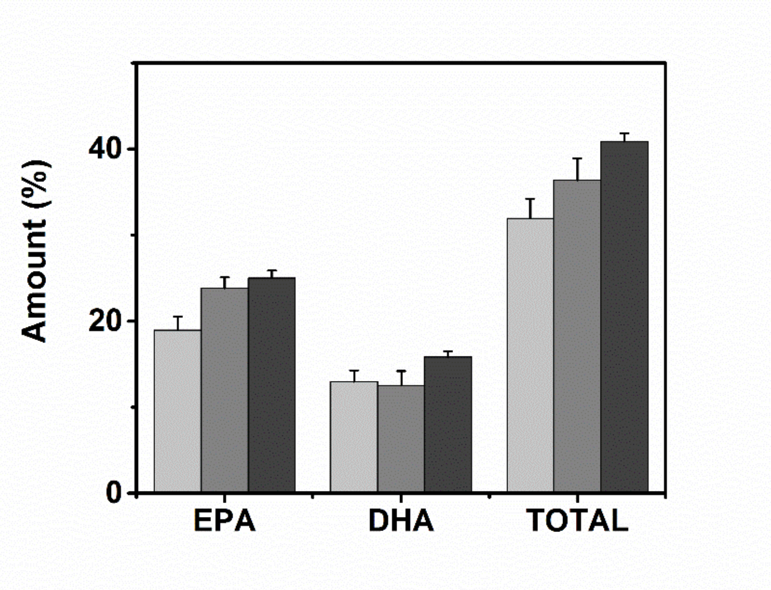


**Supplementary Figure 3**: Concentration of EPA and DHA in the glyceride fraction of unhydrolyzed anchovy oil (Light grey bars) and after 20 percent hydrolysis of anchovy oil by GTL (Dark grey bars) and DM-GTL (Black bars).

| Amino acid position | Forward primer | Reverse primer |
|---|---|---|
| 170 | TGACGGGACGACG**NNK**GTCAACATGGTTG | CAACCATGTTGAC**MNN**CGTCGTCCCGTCA |
| 171 | GGACGACGCTT**NNK**AACATGGTTGATTTC | GAAATCAACCATGTT**MNN**AAGCGTCGTCC |
| 244 | ATACCGCCCGCTACGAT**NNK**TCCGTTCCC | GGGAACGGA**MNN**ATCGTAGCGGGCGGTAT |
| 319 | GAACGACGGC**NNK**GTCAATACCATTTCGA | TCGAAATGGTATTGAC**MNN**GCCGTCGTTC |
| 358 | GTCGAC**NNK**TTGGAAATCATCGGCGTTGACCC | GGGTCAACGCCGATGATTTCCAA**MNN**GTCGAC |
| 359 | AATGTCGACCAT**NNK**GAAATCATCGGCGT | ACGCCGATGATTTC**MNN**ATGGTCGACATT |

| Mutations | Forward primer | Reverse primer |
|---|---|---|
| L170G V171R | GACGGGACGACG**GGTTGG**AACATGGTTG | CAACCATGTT**CCAACC**CGTCGTCCCGTC |
| L359C | GTCGACCATT**GTG**AAATCATCGGCG | CGCCGATGATTT**CAC**AATGGTCGAC |
| V171L | GGACGACGCTT**CTC**AACAT | ATGTT**GAG**AAGCGTCGTCC |
| L183F | GCTTTTTTGAC**TTC**ATGAAAGCG | CGCTTTCAT**GAA**GTCAAAAAAGC |

**Supplementary Table 1:** Primers used in SSM (Top) and for SDM (Bottom). Mutations are shown in bold.
